# The bone marrow microenvironment of RAS pathway mutant B-ALL is enriched for immunosuppressive regulatory T cells

**DOI:** 10.64898/2026.03.17.712388

**Authors:** Mauricio Nicolás Ferrao Blanco, Bexultan Kazybay, Alicia Perzolli, Lennart Kester, Olaf Heidenreich, Hermann Josef Vormoor

**Author notes:** Corresponding author: Hermann Josef Vormoor. < >.

## Abstract

Somatic mutations in the RAS pathway are highly prevalent in B-Cell Acute Lymphoblastic Leukemia (B-ALL), yet their impact on the bone marrow immune microenvironment and response to immunotherapy remains poorly defined. In this study, we integrated bulk RNA-sequencing, single-cell RNA-sequencing (scRNA-seq), and spectral flow cytometry to characterize the immune landscape of RAS-mutant B-ALL. We identified pathogenic mutations in KRAS, NRAS, PTPN11, or BRAF in 42% of the cohort, predominantly as clonal events. Despite similar T-cell frequencies by flow cytometry, bulk transcriptomes from RAS-mutant samples showed suppression of immune-response and T-cell-activation pathways, and T cells from RAS-mutant patients exhibited impaired proliferation ex vivo. Single-cell analysis revealed higher CD8 dysfunction scores and enrichment of regulatory T cells (Tregs) in RAS-mutant bone marrow. These findings were validated by spectral flow cytometry and by CIBERSORTx deconvolution of bulk data. Trajectory analysis supported a higher CD4 to Treg differentiation in the RAS-mutant niche, and CellChat mapping identified contact-dependent and checkpoint interactions (including TIGIT–NECTIN2 and CTLA-4–CD86/ICOSL) enriched in RAS-mutant samples. Functionally, blinatumomab produced limited leukemic-cell killing *ex vivo* overall, but addition of CTLA-4 blockade (ipilimumab) selectively restored blinatumomab efficacy in RAS-mutant samples. Together, these results indicate that RAS-pathway activation associates with a Treg-enriched, immunosuppressive bone-marrow microenvironment and point to CTLA-4–targeted strategies to enhance T-cell-engager efficacy in this subgroup.

## Introduction

B-Cell Acute Lymphoblastic Leukemia (B-ALL) is the most common pediatric malignancy, characterized by the accumulation of immature B-cell precursors in the bone marrow. While survival rates have improved significantly, relapsed and refractory disease remains a leading cause of cancer-related death in children [1]. The genomic landscape of B-ALL is well-characterized, with somatic mutations in the RAS pathway, including KRAS, NRAS, PTPN11, and BRAF, representing one of the most frequent oncogenic drivers [2–4]. Comprehensive mutational screening has demonstrated that RAS pathway alterations are present in approximately 35% of diagnostic samples and up to 25% of relapsed cases [5]. When detected at relapse, these mutations are present in the dominant clone and are associated with chemoresistance [6, 7].

Beyond their role in intrinsic tumor cell proliferation and survival, RAS pathway mutations are increasingly recognized as potent modulators of the tumor microenvironment (TME) across various malignancies [8]. In solid tumors, oncogenic RAS signaling has been shown to drive immune evasion by orchestrating an immunosuppressive landscape [9]. KRAS-mutant cells can actively recruit immunosuppressive cells and upregulate checkpoint ligands, providing a rationale for combinatorial strategies targeting both the oncogene and the immune system. This approach has been applied to colorectal cancer, combined inhibition of BRAF, MEK, and PD-1 has shown promise by augmenting the tumor immune response [10]. Similarly, in lung cancer models, KRAS G12C inhibition has been found to drive anti-tumor immunity and synergize with immune checkpoint blockade, transforming a “cold” immune microenvironment into a pro-inflammatory state [11].

Despite this growing body of evidence in solid tumors, the immunomodulatory consequences of RAS pathway mutations in the specific context of B-ALL remain poorly defined. While the genetic prevalence of these mutations is established, it is currently unknown whether RAS-mutant lymphoblasts actively shape the bone marrow immune niche to favor immune escape. This knowledge gap needs to be addressed, given the shift in B-ALL treatment from chemotherapy towards immunotherapy. The bispecific T-cell engager blinatumomab has significantly improved disease-free survival in patients with B-ALL [12–18]. However, resistance to blinatumomab occurs, and efficacy is mostly reliant on the quality and fitness of the patient’s T cell compartment [19].

Understanding the interplay between RAS mutations and immune effector cells may, therefore, be important for optimizing immunotherapy in B-ALL. In this study, we aimed to dissect the immune landscape of RAS-mutant B-ALL using single-cell transcriptomics and functional assays. We hypothesized that RAS pathway alterations drive a specific immune escape phenotype that can be therapeutically targeted. Here, we report that RAS mutations are associated with a Treg-enriched microenvironment and indicate that targeting this suppression via CTLA-4 blockade may restore the efficacy of blinatumomab.

## Results

### RAS pathway mutations characterize a distinct immunomodulatory landscape in B-ALL

To characterize the impact of RAS pathway alterations, we analyzed bulk RNA-seq data from a cohort of 210 samples derived from bone marrow mononuclear cells (BMMC) of patients with B-ALL. Mutation calling identified pathogenic alterations in KRAS, NRAS, PTPN11, and BRAF in 42% (87/210) of cases (Figure 1A). Specific variants identified included common hotspots such as KRAS G12/G13, NRAS G12/G13/G61, and PTPN11 A72V/G69L, alongside others (detailed in Supplementary Table 1). Co-mutations in multiple RAS pathway genes were observed. Variant Allele Frequency (VAF) analysis revealed that most RAS pathway mutations (62/87) were in the major clone (VAF > 25%) (Figure 1B). Consistent with previous reports [4], we observed a high incidence of RAS pathway mutations in ZNF384-rearranged and hyperdiploid subtypes, whereas incidence was lower in ETV6::RUNX1 and absent in BCR::ABL1 ALL (Figure 1C). To assess the transcriptomic consequences of these alterations, we focused our differential expression analysis on samples harboring high allele frequency mutations (VAF > 25%; n = 62) compared to wild-type counterparts (n = 123) (Supplementary Table 2). Gene ontology analysis revealed a significant suppression of pathways related to immune response and T cell activation in RAS-mutant samples (Figure 1D & Supplementary Table 3), which correlated with decreased expression of cytolytic and effector molecules, including *GZMM*, *GZMK*, and *IFNG* (Figure 1E). We next investigated whether this transcriptomic suppression reflected a quantitative reduction in T cells. Flow cytometric analysis of diagnostic BMMC samples (n = 74 mutant; n = 116 wild-type) showed no significant difference in the percentage of T cells (CD45+CD3+) between groups (Figure 1F). However, functional assays revealed a qualitative defect. Upon *ex vivo* stimulation with anti-CD3/CD28 beads, T cells from RAS-mutant patients (n = 7) failed to proliferate, exhibiting significantly lower CFSE dilution compared to wild-type samples (n = 7) (Figure 1G-H). These data suggest that while RAS pathway mutations do not alter T cell abundance, they are associated with profound T cell hypoactivation.

**Figure 1.**
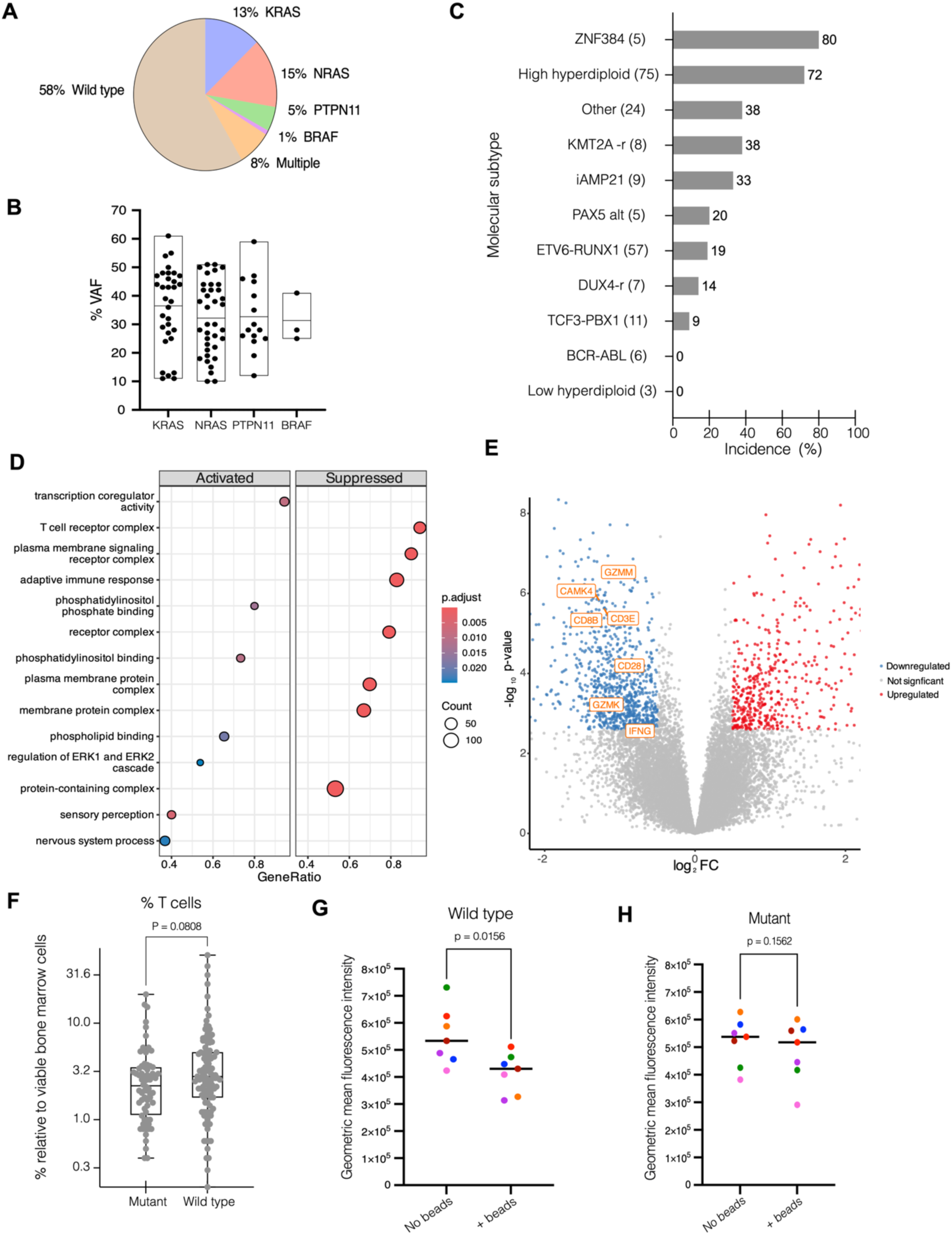
RAS pathway mutation and immune phenotype in B-ALL. A) Distribution of RAS pathway mutations in the cohort. “Multiple” indicates samples with two or more mutated RAS pathway genes. B) Variant allele frequency (VAF) for RAS pathway mutations. Each dot represents one sample. For samples with co-mutations, the highest VAF for the respective gene is shown. C) Frequency of RAS pathway mutations across molecular subtypes. In parentheses the number of samples. D) Gene ontology representation of the top significantly activated and suppressed molecular processes in RAS mutant samples. E) Volcano plot of differentially expressed genes comparing clonal RAS mutant (blue) versus wild type (red) samples. T cell cytolytic and effector genes are labeled. F) Percentage of T cells (CD45+CD3+) in diagnostic bone marrow mononuclear cells determined by flow cytometry (n = 74 mutant; n = 116 wildtype. Statistical comparison by unpaired two-tailed t-test. G, H) T cell proliferation measured by CFSE geometric mean fluorescence intensity (gMFI) in CD3+ T cells at day 5, with or without anti-CD3/CD28 stimulation, for 7 wild type and 7 RAS mutant patient samples. Each dot represents the mean gMFI of technical triplicates; every color denotes a different patient. Statistical analysis by two-tailed Wilcoxon matched-pairs test.

### Single-cell transcriptomics reveals T cell dysfunction in the RAS mutant B-ALL bone marrow microenvironment

To dissect the cellular heterogeneity underlying this immune suppression, we performed single-cell RNA sequencing (scRNA-seq) on bone marrow mononuclear cells from 8 patients (Figure 2A). We employed flow cytometry to isolate leukemic plus normal B-lineage cells (CD19+), non-leukemic hematopoietic cells (CD19-CD45+ and CD19-CD235a+), and non-hematopoietic cells (CD19-CD45-CD235a-), an approach used in our previous study [20]. Following quality control, 19,199 cells were analyzed and annotated into 14 distinct populations (Figure 2B, Supplementary Figure 1). Patient contributions to clusters were variable overall, but contributions to the T cell cluster were similar across patients (Figure 2C). Sub-clustering of the T cell compartment identified five major clusters: CD4+ T cells, CD8+ T cells, Gamma-delta T cells, Mucosal-associated invariant T (MAIT) cells, and Tregs (Figure 2D, Supplementary Figure 2). These cell types were characterized by canonical marker expression (Figure 2E, Supplementary Table 4). To test whether T cells in the RAS mutant microenvironment display an immunosuppressive phenotype, we computed a CD8 dysfunction score derived from inhibitory checkpoint and exhaustion-associated genes, including HAVCR2 (TIM3), TIGIT, LAG3 and the transcription factor TOX [21–23]. CD8+ T cells from RAS-mutant bone marrow exhibited significantly higher dysfunction scores than those from wild-type samples (Figure 2F), corroborating the hypoactivation observed in bulk assays.

**Figure 2.**
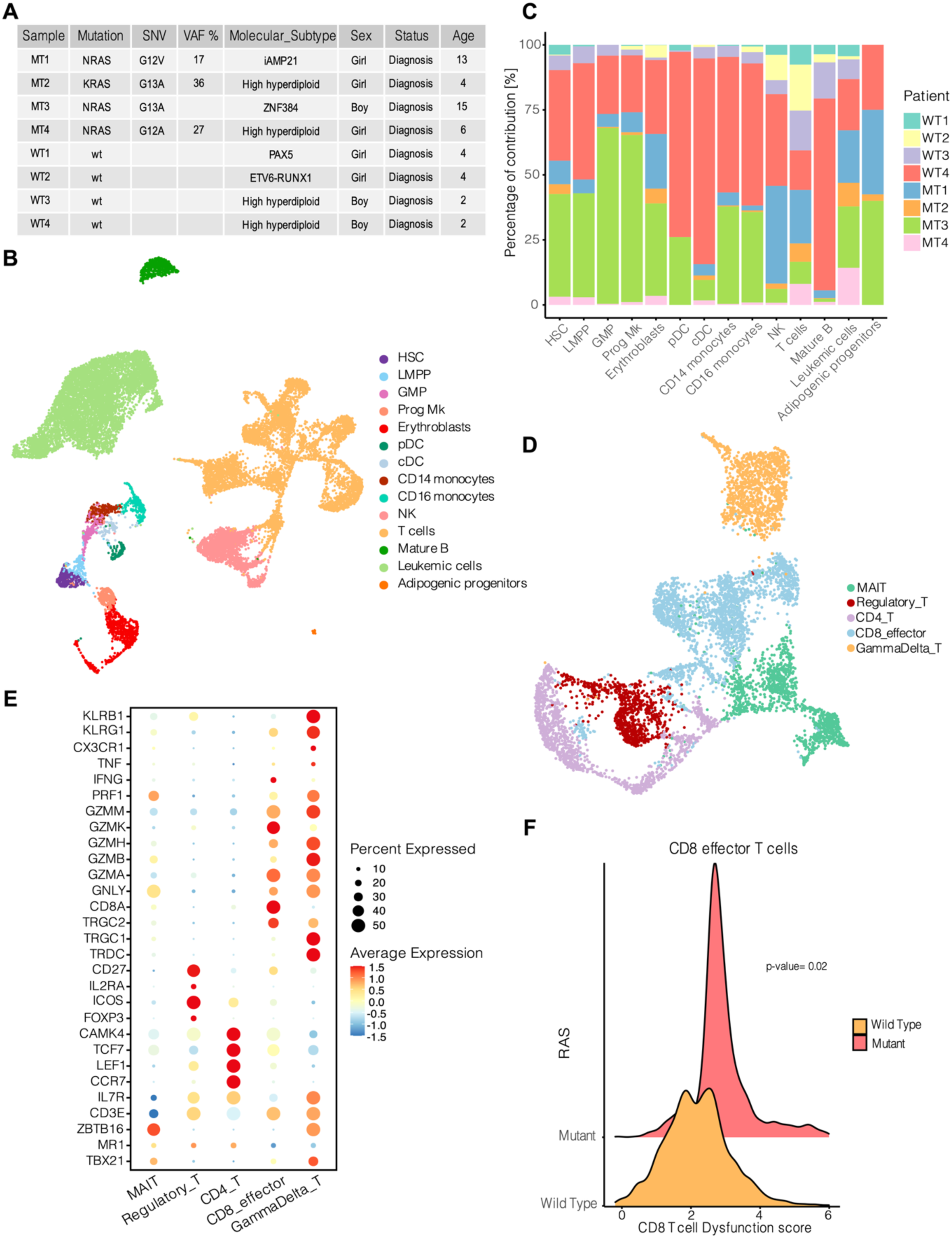
Single-cell profiling of the immune bone marrow niche in B-ALL. A) UMAP of 19,199 high-quality bone marrow mononuclear cells from 8 patients, annotated into 14 major cell populations (labels shown). B) Table indicating patient samples characteristics. C) Stacked bar plot showing the patient-wise contribution to each annotated cluster (each bar = one cluster; colors denote patients). D) UMAP of the T cell compartment after sub-clustering, showing five discrete T cell subsets: CD4+ T cells, CD8+ T cells, gamma-delta (γδ) T cells, MAIT cells, and regulatory T cells (Tregs). E) Dotplot of canonical marker genes used to define T cell subsets. F) Ridge plot of CD8 T cell dysfunction score in CD8+ T cells, computed from inhibitory and exhaustion-associated genes (including HAVCR2/TIM3, TIGIT, LAG3, TOX). Each distribution shows single-cell dysfunction scores from CD8 effector T cells pooled by RAS status. For statistical testing, single-cell scores were summarized to patient-level pseudo-bulk means (n = 4 RAS-mutant, 4 wild-type) and a two-sided Wilcoxon rank-sum test was performed.

### Regulatory T cells are enriched in RAS-mutant B-ALL

Comparison of cell type proportions within the scRNA-seq data indicated an abundance of Tregs in RAS-mutant samples (Figure 3A, B). Transcriptomic analysis further showed an increase of checkpoint molecules in Tregs in the RAS-mutant niche, namely CTLA4 and TIGIT (Figure 3C, Supplementary Figure 2, Supplementary Table 5). These immune checkpoints are key regulators of excessive T cell activity and can be exploited by tumors to suppress anti-tumor immunity [24]. We validated the scRNAseq finding by spectral flow cytometry on an independent cohort of diagnostic BMMC samples (n = 12; 6 mutant, 6 wild-type). To reduce subtype confounding we restricted this comparison to hyperdiploid cases. Frequencies of total T cells, leukemic cells, CD4+, CD8+, and gamma delta T cells were comparable between groups (Figure 3D & Supplementary Figure 3), whereas Tregs were significantly enriched in RAS-mutant samples (Figure 3E). The percentage and median fluorescence intensity (MFI) of checkpoint/ exhaustion markers (TIM3, LAG3, TIGIT, PD-1, CTLA-4) on Tregs and CD8+ T cells did not differ significantly between groups (Supplementary Figure 3), suggesting that the immunosuppressive phenotype may be driven by Treg abundance rather than per cell upregulation of checkpoint molecules. To generalize these findings, we performed cell type deconvolution (CIBERSORTx) on the larger bulk RNA-seq cohort (n = 185). This analysis confirmed significantly increased Treg levels in RAS-mutant patients (Figure 3F). Importantly, this enrichment was observed across all mutated genes and remained consistent when analyzing hyperdiploid samples separately or combining other subtypes (Supplementary Figure 4). Notably, some RAS wild-type samples also exhibited high inferred Treg levels; because BCR::ABL activates MAPK/ERK signaling similarly to RAS mutations, we compared Treg estimates in BCR::ABL(+/-like) versus other wild-type samples and observed increased Tregs in the BCR::ABL(+/-like) group (Figure 3G). We also observed a positive association between inferred Treg abundance and ERK/MAPK pathway activity (Supplementary Fig. 4). Together, these results indicate that Treg enrichment is associated with RAS/MAPK pathway activation, independent of broader molecular subtype.

**Figure 3.**
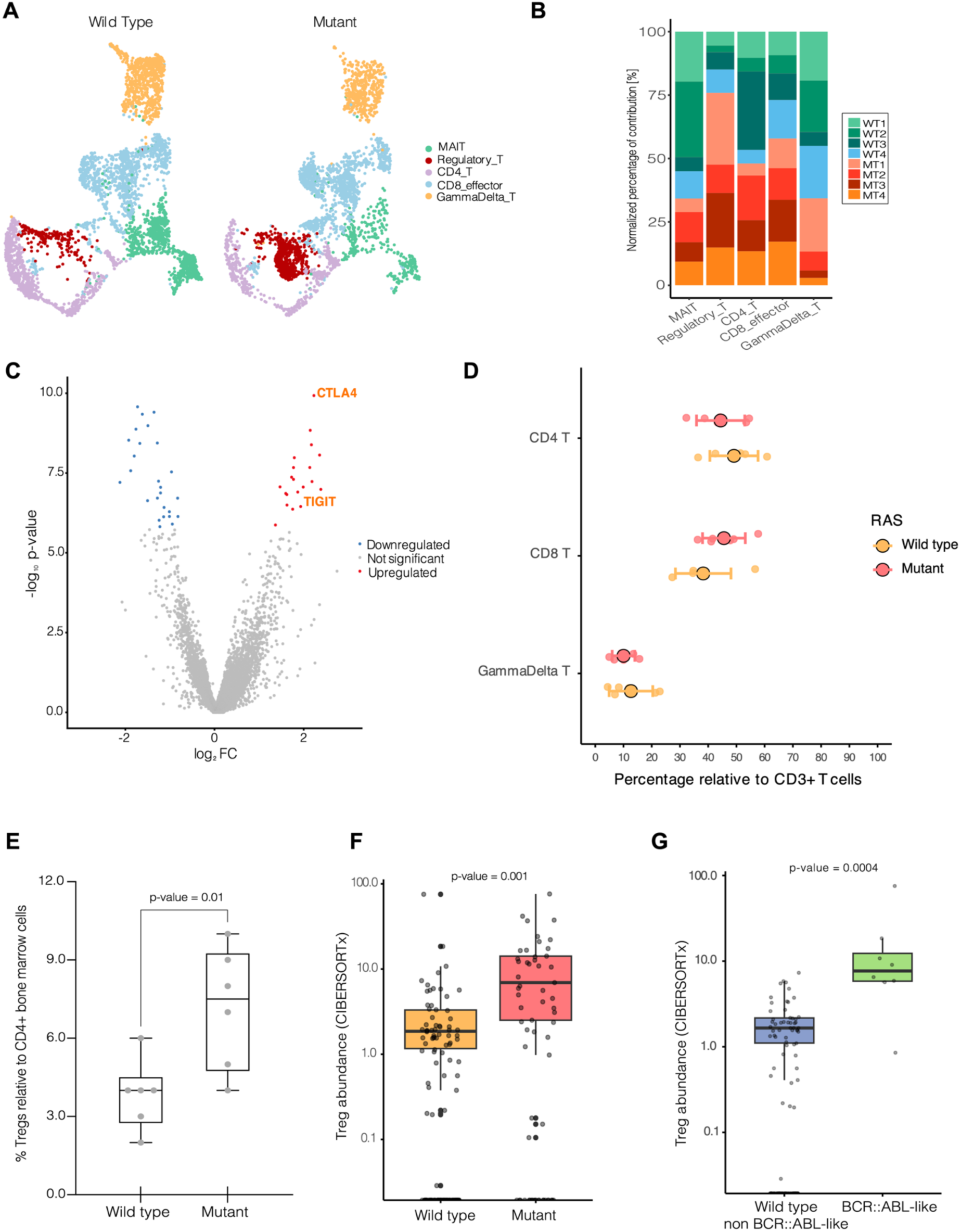
Regulatory T cell enrichment in RAS-mutant B-ALL. A) UMAPs of the T cell compartment from scRNA-seq, displayed side-by-side: right, cells from RAS-mutant samples; left, cells from wild-type samples. Major T cell clusters are labeled. B) Stacked bar plot showing the patient-wise contribution to each annotated T cell cluster; each bar corresponds to one cluster and colors denote individual patients, demonstrating greater relative Treg contribution from RAS-mutant samples (p-value = 0.02). C) Volcano plot of differentially expressed genes in Tregs comparing the ones in the RAS mutant niche (red) versus wild type (blue) samples. D) Spectral flow cytometry analysis of T cell populations in bone marrow cells in an independent cohort (n = 12; 6 RAS-mutant, 6 wild-type samples). Normality was assessed with Shapiro–Wilk and variances with Levene’s test. Group comparisons (RAS-mutant vs wild-type) were performed per cell type using Welch’s two-sample t-test with p-values adjusted for multiple testing by the Benjamini–Hochberg procedure. No comparisons were significant after adjustment. E) Box-and-whisker plot of Treg frequency relative to CD4+ bone marrow cells, same samples as in panel D. Tregs were defined as (CD4+CD25+CD127lowFOXP3+). Boxes show median and interquartile range. Statistical comparison performed using a two-tailed unpaired t-test. F) CIBERSORTx impute cell fractions of Tregs on bulk RNA-seq (n = 185) showing regulatory T cell estimates for each sample. Absolute scores reflect relative estimated cell abundance across samples and are not direct cell percentages. Group comparison (RAS-mutant vs wild-type) was performed using a two-sided Wilcoxon rank-sum test. G) Boxplot of CIBERSORTx absolute Treg scores (log10 scale) comparing wild-type (WT) and BCR::ABL-like samples. Boxes show median and interquartile range; points are individual samples (jittered). Group comparison was performed with a two-sided Wilcoxon rank-sum test.

### CD4+ T-cell Differentiation and RAS-Mutant-Specific Immunosuppressive Signaling

To investigate mechanisms driving Treg expansion in RAS-mutant B-ALL, we performed trajectory analysis on the CD4-lineage subset (CD4 T cells and regulatory T cells) (Figure 4 A & B). Expression dynamics across pseudotime showed naive markers (CCR7, LEF1, TCF7) enriched early, intermediate activation/transition markers (STAT5A, CD69, TGFB1) peaking mid-trajectory, and Treg-associated genes (IL2RA, FOXP3) elevated at late pseudotime (Figure 4C). Pseudotime ordering revealed that CD4-lineage cells from RAS-mutant samples occupy more advanced pseudotime states than those from wild-type samples, consistent with increased differentiation (Figure 4 D & E). We next mapped differential expression onto an inferred ligand–receptor network to identify interactions enriched in the RAS-mutant niche (Figure 4F; Supplementary Table 6). Focusing on signals targeting CD4+ T cells and Tregs (given inducible Treg differentiation), we found that increased signaling in the RAS-mutant microenvironment originated mainly from leukemic cells, conventional dendritic cells, and CD14+ and CD16+ monocytes. Many enriched interactions were contact-dependent; notably, PECAM1-mediated interactions were detected only in the RAS-mutant niche and have been linked to reduced T-cell activation [25]. We also observed checkpoint-related ligand–receptor pairs, including TIGIT–NECTIN2, and CTLA-4 engagement with CD86 and ICOSL specifically in the RAS-mutant environment, consistent with a microenvironment that promotes Treg activity and suppresses CD8+ effector function.

**Figure 4.**
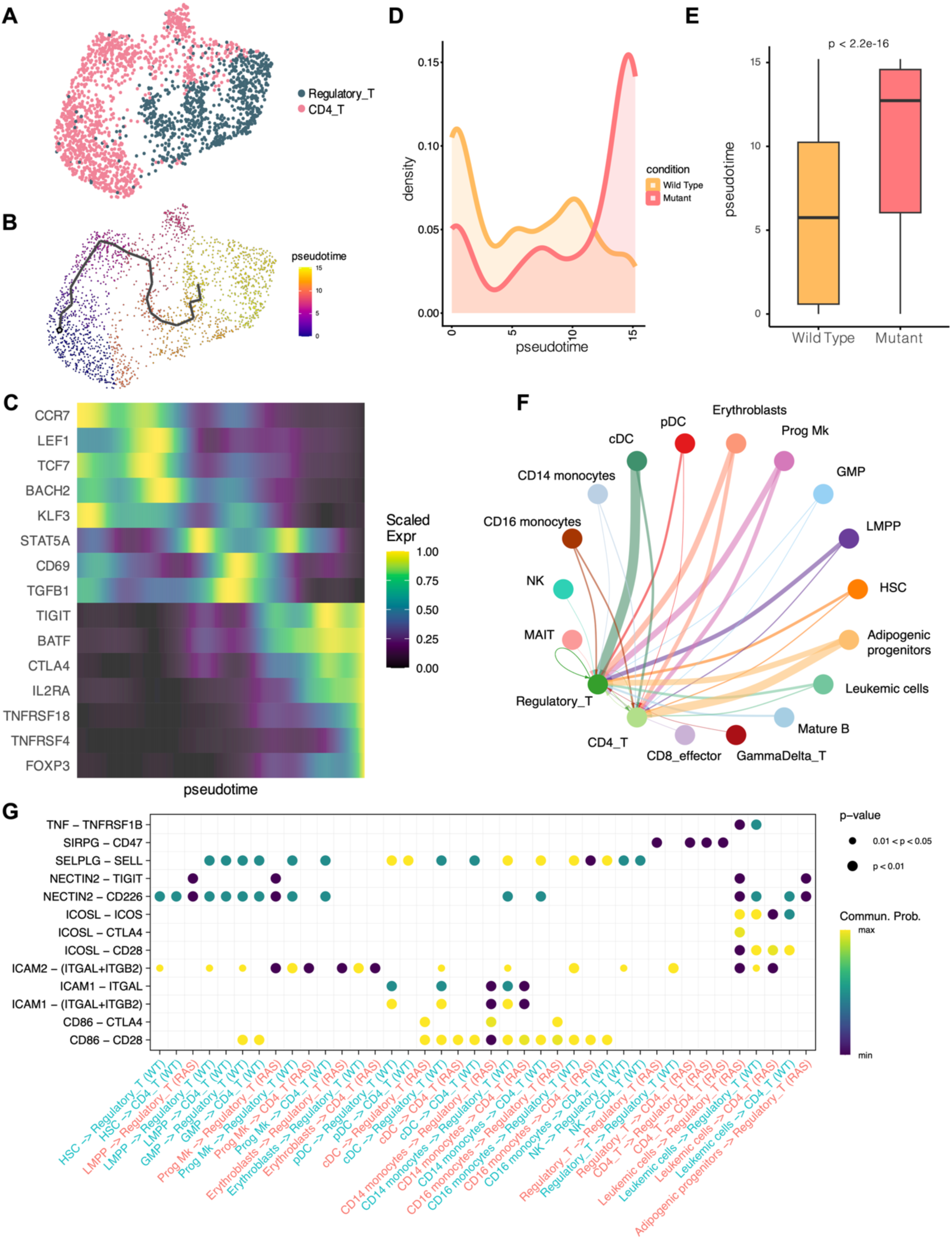
CD4+ T-cell trajectory and cell–cell communication analysis. A) UMAP of CD4+ T-cell compartment (CD4 T and regulatory T cells) from the integrated single-cell dataset. B) Monocle3 trajectory inferred from the CD4+ T-cell Seurat subset. The plotted trajectory (black curve) is overlaid on the UMAP and cells are colored by the inferred pseudotime. C) Heatmap of selected genes ordered by Monocle3-inferred pseudotime (cells ordered left to right by increasing pseudotime). Rows are genes associated with naive, activation/transition, effector/suppressive and Treg identity. Per-gene expression was extracted from the Seurat RNA assay, smoothed across ordered cells using a cubic smoothing spline (spar = 0.9), then rescaled to the 0–1 range for visualization. D) Density plot of Monocle3-inferred pseudotime for CD4+ T cells, stratified by RAS status. Curves show the kernel density of pseudotime values per condition (area normalized). E) Boxplot comparing pseudotime distributions between RAS-mutant and wild-type cells; boxes indicate median and interquartile range, whiskers extend to 1.5× IQR. Statistical comparison between groups was performed on cell-level pseudotime values using a two-sided Student’s t-test. F) Aggregate circle plot of signaling pathways upregulated in the RAS-mutant condition. Differential expression of ligand/receptor components was performed using CellChat’s DE workflow (presto/Wilcoxon; thresh.pc = 0.1, thresh.fc = 0.05, thresh.p = 0.05). Differentially expressed ligand–receptor pairs with upregulated ligands in RAS were mapped to inferred intercellular communications and prioritized pathways were selected. The circle plot displays aggregated RAS-enriched signaling to CD4_T and Regulatory_T cells. Edge thickness and arrowheads indicate interaction strength and directionality. G) Bubble plot of selected ligand–receptor pathways enriched in the RAS-mutant niche. Sources shown are the top sender cell types identified from the differential mapping; targets are T-cell populations (CD4_T, Regulatory_T). Bubble size encodes interaction strength (communication probability) and bubble color (viridis) encodes the direction/magnitude of differential signaling between RAS and WT. An expanded list of the interactions and statistics are provided in Supplementary Table 6.

### CTLA-4 blockade restores Blinatumomab efficacy in RAS-mutant B-ALL

High Treg frequency has been reported to impair clinical responses to the bispecific T cell-engager blinatumomab [26]. Because we observed Treg enrichment in RAS-mutant samples, we investigated the efficacy of blinatumomab *ex vivo.* To minimize subtype confounding, this analysis was restricted to hyperdiploid cases. Samples derive from the same patients shown in Figure 3, which have comparable leukemic-cell and CD8+ T-cell frequencies but higher Treg abundance in RAS-mutant cases. In liquid cultures of patient BMMCs, blinatumomab monotherapy resulted in low leukemic cell killing in both RAS mutant and wild type groups (Figure 5A), indicative of a broadly immunosuppressive microenvironment, in line with recent findings [27]. In RAS mutant samples, blinatumomab efficacy (% leukemic cell killing) inversely correlated with the frequency of regulatory T cells, independent of leukemic cell and total T cell percentages (Figure 5B). Given that Tregs suppress in part via CTLA-4 [28], and that CTLA-4-related interactions were enriched in the RAS mutant niche inferred by our cell-cell communication analysis, we tested whether CTLA-4 blockade could enhance blinatumomab activity. Addition of ipilimumab to blinatumomab had a significant increase in leukemic cell killing in RAS-mutant samples, while no significant effect in wild-type samples was observed (Figure 5C, D). These data indicate that Treg-mediated immune evasion contributes to blinatumomab resistance in RAS-mutant B-ALL, and that combining CTLA-4 blockade with blinatumomab may help overcome this resistance.

**Figure 5.**
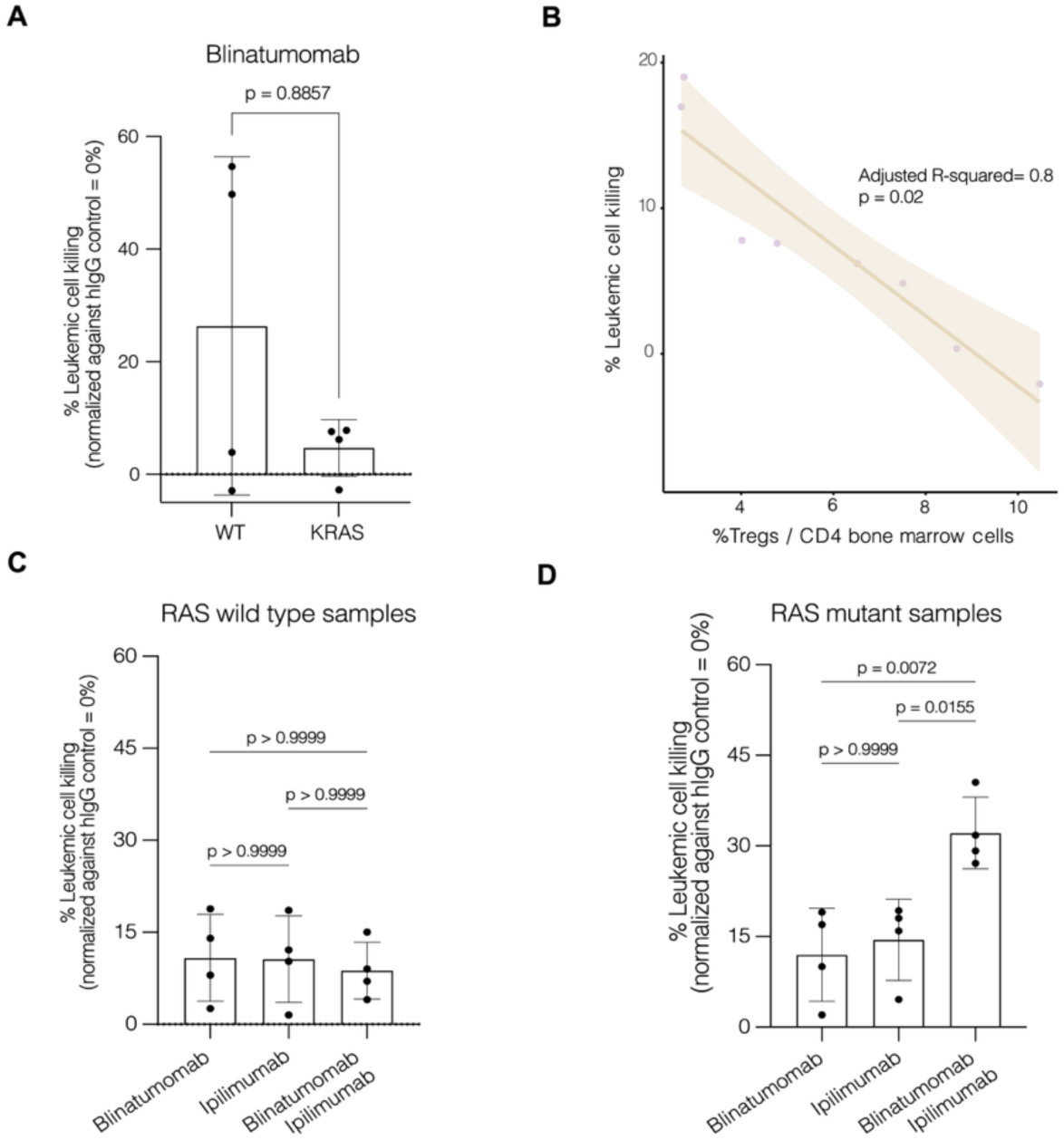
Ipilimumab Enhances Blinatumomab Killing in RAS-Mutant B-ALL. A) Ex vivo leukemic-cell killing by blinatumomab monotherapy in liquid cultures of patient BMMCs (n = 8 total; 4 RAS-mutant, 4 wild-type). Statistical analysis was performed using a two-tailed Mann–Whitney U test. B) Relationship between regulatory T cell frequency and ex vivo blinatumomab response in RAS mutant samples (n=8). Scatterplot shows sample-level blinatumomab response (% leukemic cell killing) versus % Tregs among CD4+ bone marrow cells with fitted regression line and 95 % CI. Spearman correlation: rho= 0.976, p= 3.97x10-4. Multivariable linear regression analysis (Blina_response ∼ Treg_CD4 + Leukemic_cells + T_cells) indicates that the % Tregs is independently associated with lower blinatumomab response, while leukemic cells and T cells were not significant. C, D) Leukemic-cell killing following treatment with blinatumomab alone, ipilimumab alone or blinatumomab + ipilimumab, shown separately for RAS-mutant and wild-type patient samples (4 RAS-mutant, 4 wild-type). Differences across treatment groups were tested by one-way ANOVA with Bonferroni multiple-comparison post-hoc correction. In all graphs the bars show group means; error bars indicate ± SD. Individual patient values are overlaid as points. Leukemic-cell killing was calculated as percent reduction in viable leukemic cells relative to IgG controls.

## Discussion

In this study, we define a distinct immunomodulatory role for RAS pathway mutations in B-Cell Acute Lymphoblastic Leukemia (B-ALL). While RAS mutations have long been recognized as drivers of leukemic proliferation and chemoresistance in relapsed ALL [2, 5], our data reveal that they also actively orchestrate a profoundly immunosuppressive bone marrow microenvironment. By integrating bulk and single-cell transcriptomics with functional assays, we demonstrate that RAS-mutant B-ALL is characterized by intrinsic T cell dysfunction and a specific enrichment of Tregs, creating a barrier to immune surveillance that can be overcome by combinatorial checkpoint blockade.

Our findings in B-ALL parallel the “immune-cold” phenotypes observed in RAS-driven solid tumors [29]. Just as KRAS mutations in lung and colorectal cancers drive immune evasion through the recruitment of suppressive cells and upregulation of checkpoints [9, 11], we show that RAS-mutant lymphoblasts condition the bone marrow niche to disarm anti-leukemic immunity. Specifically, while flow cytometry showed no quantitative reduction in total T cell numbers, scRNA-seq revealed a qualitative defect, with T cells in the RAS-mutant niche exhibiting high dysfunction scores and a failure to proliferate upon stimulation. This suggests that the resistance mechanisms described in solid tumors, where RAS signaling facilitates crosstalk with the microenvironment to drive malignancy [30], are conserved in hematologic malignancies, extending the relevance of RAS as an immunomodulatory oncogene to leukemia.

A central finding of this work is the specific enrichment of Tregs in RAS-mutant patients, independent of the broader molecular subtype (e.g., Hyperdiploid). This robust association provides a biological mechanism for the observed T cell hypoactivation. Tregs exert suppression via contact-dependent mechanisms, prominently including the constitutive expression of CTLA-4, which downregulates costimulatory signaling [28, 31]. The presence of this high Treg burden is clinically significant given the widespread adoption of blinatumomab in B-ALL protocols [32, 33]. Previous studies have indicated that high Treg frequency correlates with blinatumomab non-response [26]. Consistent with this, our ex vivo assays demonstrated that blinatumomab monotherapy had limited efficacy in the RAS-mutant context, likely due to the suppressive constraints imposed by the Treg-enriched environment.

Crucially, however, we demonstrate that this immune resistance is reversible. The addition of ipilimumab, a CTLA-4 inhibitor, significantly enhanced blinatumomab-mediated leukemic cell killing specifically in RAS-mutant samples. This aligns with recent successes in BRAF-mutant colorectal cancer, where combining MAPK pathway inhibition with immune checkpoint blockade improved responses [10]. In our context, targeting the specific mechanism of suppression (CTLA-4 on Tregs) restored the cytotoxic potential of T cells engaged by blinatumomab. This suggests that in RAS-mutant B-ALL, CTLA-4 acts as a critical, non-redundant checkpoint maintaining immune tolerance.

The observation that RAS-mutant lymphoblasts reside within a Treg-enriched, immunosuppressive niche provides a compelling rationale for combinatorial strategies that target both the oncogenic driver and the immune system. While specific inhibitors targeting the KRAS G12C mutation have revolutionized treatment in solid tumors [34, 35], this specific variant has a low incidence in B-ALL; indeed, only 1 of the 87 RAS-mutant patients in our cohort harbored the G12C mutation. Consequently, given the lack of specific inhibitors for the broad spectrum of RAS variants found in B-ALL, downstream inhibition of the MAPK/ERK pathway remains the primary therapeutic avenue. MEK inhibitors, such as selumetinib, have demonstrated efficacy ex vivo and in clinical settings for RAS-mutant ALL [7, 36, 37]. Therefore, combining selumetinib with blinatumomab represents a promising therapeutic approach.

Our study has limitations, including the use of *ex vivo* models which may not fully recapitulate the systemic pharmacodynamics of checkpoint blockade, and the need for validation in larger clinical cohorts to confirm Tregs and RAS mutations as predictive biomarkers for antibody-based and/or cellular immunotherapies. Furthermore, while we identified the association between RAS mutations and Treg abundance, the precise molecular signals secreted by RAS-mutant blasts to recruit these cells remain to be fully elucidated.

In conclusion, we identify RAS pathway mutations as a potential biomarker for a Treg-enriched, immunosuppressive microenvironment in B-ALL. These data extend the view of RAS mutations solely as proliferation drivers and support a precision medicine approach. For patients with RAS-mutant B-ALL, therapeutic strategies combining T cell engagers with CTLA-4 blockade may offer a mechanism to dismantle the immunosuppressive niche and unleash effective anti-leukemic immunity.

## Materials and methods

### Bone marrow samples

Bone marrow aspirates were obtained from pediatric B-ALL patients in conjunction with routine clinical sampling after obtaining informed consent from parents and/or legal guardians in accordance with institutional review board approval at the Princess Máxima Center (application number PMCLAB2022.305). Bone marrow mononuclear cells (BM-MNCs) were isolated from fresh bone marrow aspirates using density gradient centrifugation and cryopreserved in liquid nitrogen.

### Differential expression analysis on bulk RNA seq samples

Raw gene counts (n = 185 samples) and sample metadata (molecular subtype, RAS status, VAF, SNV, Clone) were imported into R. Factors were set appropriately and DESeq2 was used to test for differential expression between RAS-mutant and wild-type samples with the design formula ∼ Molecular_subtype + RAS to account for molecular subtype effects [38]. DESeq was run on the count matrix and results for the RAS contrast (mut vs wt) were extracted. Adjusted p-values (Benjamini–Hochberg) were used. Significant differentially expressed genes (DEGs) were defined as padj < 0.05 and absolute log2 fold change > 0.5. For volcano visualization, a base mean filter (baseMean > 200) was applied to reduce noise.

Gene-set enrichment analysis was performed using clusterProfiler. A ranked gene list of log2 fold-changes (named by gene symbol) from the filtered DEG set was supplied to gseGO (OrgDb = org.Hs.eg.db, keyType = “SYMBOL”, minGSSize = 10, maxGSSize = 1000, pvalueCutoff = 0.05). Results were visualized with dotplot and gseaplot.

### FACs enrichment of stromal, hematopoietic and leukemic cells for scRNAseq

Cryopreserved bone marrow mononuclear cells were thawed and resuspended in RPMI 1640 medium containing 5% FBS and 1 mM EDTA. The cell suspension was passed through a 70 μm mesh and stained for 20 minutes at 4 degrees with an antibody cocktail containing CD19-PE (Catalogue 302207, Clone HIB19, 1:100), CD235a-FITC (Catalogue 349103, Clone HI264, 1:100), CD45-PE-Cy7 (Catalogue 304015, Clone HI30, 1:100), all from Biolegend. Cells were washed twice with medium followed by 7AAD staining (Catalogue 420404, 1:100) to exclude dead cells before sorting on a Sony SH800S instrument (Sony Biotechnology). Leukemic cells, hematopoietic, and non-hematopoietic cells were sorted in RPMI 1640 medium containing 5% FBS. Samples were kept during the whole procedure at 4 °C. Following enrichment, each of the three cell fractions (leukemic, hematopoietic, non-hematopoietic) were resuspended at a concentration of 1000 cells/μL in PBS + 0.04% BSA. Cells were pooled in a 1:1:1 ratio and the cell suspension was subsequently mixed with a cell suspension from another patient with a different gender (multiplexed boy/girl samples). Subsequently, the cell suspension was loaded into the 10x Genomics Chip as described in ‘Single-cell RNA library generation’.

### Single-cell RNA-seq library generation

In total, 20,000 cells were loaded for each (multiplexed) sample onto a 10x Genomics Chip G. We used Chromium Next GEM Single Cell 3’ GEM, Library & Gel Bead Kit and followed the protocol as provided by 10x Genomics. Sequencing was performed on Illumina NovaSeq 6000 sequencer and parameters were set according to 10x Genomics recommendations.

### Data processing and filtering

Reads from the sequencing were aligned to the human reference genome (GRCh38) using Cell Ranger v7.1.0. Samples were demultiplexed using souporcell based on Single Nucleotide Polymorphisms (SNPs) identified within the reads. Patient identities were assigned based on specific sex chromosome genes. Further analysis was performed using R v4.0.2 (https://www.r-project.org/) including several packages which are mentioned in the according section. As an initial step, we filtered out low-quality cells. Cells with less than 200 expressed features were removed. Additionally, cells with greater than 10% of reads mapping to mitochondrial genes were excluded. Doublets/multiplets, which can arise during library preparation, were identified and removed using the scDblFinder package with default parameters [39].

### Data integration

All patient scRNAseq datasets were merged into one Seurat object after prefiltering (see section: “Data processing and filtering”). Based on this object, the data was normalized using the SCTransform function from Seurat with default parameters. Principal component analysis (PCA) was performed using the RunPCA function from Seurat, with the number of principal components (npcs) set to 30, a value determined empirically to capture the major sources of variation in the data. The IntegrateLayers function was run with the RPCA integration method and default settings to integrate the data across different patients and account for batch effects.

### Annotation

Cluster annotation for the full integration of all samples was performed at a low-resolution to identify major cell types. Cell types were annotated based on information from literature, functional information from gene sets based on the marker genes of a cluster and unbiased cell type recognition using the Human Primary Cell Atlas and bone marrow scRNAseq references [20, 40, 41] in SingleR with default parameters [42].

### Marker gene identification

Marker genes per cluster were calculated by the Seurat function FindAllMarkers, using the method “Model-based Analysis of Single-cell Transcriptomics” (MAST) with a log fold change threshold of 0.3 and min-pct set to 0.3.

### CD8 T cell dysfunction score

A CD8 T-cell dysfunction score was computed on the Seurat object using a four-gene signature (HAVCR2, TIGIT, LAG3, TOX). The default assay was set to “RNA” and module scores were generated with AddModuleScore, yielding the CD8TcellDysfunction. Scores for CD8 effector cells were visualized as a ridge plot grouped by RAS status. For statistical testing, per-cell scores were extracted with Patient_ID and RAS status, then averaged to patient-level pseudo-bulk means (one mean per patient) using group_by (Patient_ID, RAS) and summarise (Mean_Score = mean (CD8TcellDysfunction)), yielding n = 8 patient means (4 RAS-mutant, 4 wild-type). Group comparisons were performed on these patient means using a two-sided Wilcoxon rank-sum test (nonparametric). The Wilcoxon W and p-value are reported in the figure.

### Monocle3 trajectory and pseudotime analysis

Seurat CD4+ T-cell subset objects were converted to Monocle3 cell_data_set objects for trajectory inference. Raw count layers for individual patient batches were extracted from the Seurat RNA assay and combined into a single counts matrix (genes × cells) that matched the Seurat cell ordering. Cell metadata and gene annotation (gene_short_name = SYMBOL) were transferred to the Monocle3 object. The RPCA UMAP coordinates computed in Seurat (reduction = “umap.rpca_CD4T”) were assigned to reducedDims(cds)$UMAP in Monocle3 so that Monocle3 used the same embedding for graph learning. Cells were clustered with cluster_cells(cds, reduction_method = “UMAP”), the principal graph was learned using learn_graph(cds, use_partition = TRUE), and cells were ordered along pseudotime with order_cells(cds). Pseudotime values were stored in the cell metadata and used for downstream comparisons. To identify genes whose expression varies along the trajectory we applied Monocle3’s graph_test (Moran’s I spatial autocorrelation) on the principal_graph neighborhood (neighbor_graph = “principal_graph”); genes with q < 0.05 and high Moran’s I were prioritized. For heatmap visualization, we selected biologically relevant genes (naive markers, early activation/transition, effector/suppressive and Treg identity genes), extracted normalized expression from the Seurat RNA assay (FetchData), ordered cells by increasing pseudotime, smoothed each gene’s expression across ordered cells using smoothing splines (spar = 0.9), and rescaled each gene to the 0–1 range for display. Heatmaps and density/box plots of pseudotime distributions were generated with ggplot2.

### Cell-cell communication analysis

The CellChat algorithm was applied to perform an unbiased ligand-receptor interaction analysis [43]. CellChat’s human ligand-receptor database was employed to infer potential cell-cell interactions. Overexpressed genes and interactions were identified, followed by projection onto a human protein-protein interaction network. Communication probabilities were computed using a tri-mean method, and signaling pathways were aggregated and filtered to include interactions supported by at least 10 cells per group. Network centrality metrics were calculated, and interaction patterns were visualized via circle plots and bubble plots.

### Inference of copy number variation

To distinguish leukemic cells from non-malignant populations in pediatric B-ALL samples, we performed copy number alteration (CNA) analysis on individual patients using the SCEVAN (Single-Cell Evolutionary Variational Autoencoder for CNAs) R package [44]. For each patient, we extracted single-cell gene expression data from an integrated Seurat object. To define a set of reference “normal” cells for CNA inference, clusters corresponding to T cells and monocytes were identified based on expression of canonical markers such as CD3G, CD8A, CD14, and FCGR3A, using DotPlot() visualization. Cells from clusters expressing these markers were selected as the normal reference population. Gene expression matrices were extracted from the RNA assay for each sample and converted to matrix format for input into the pipelineCNA() function from the SCEVAN package. SCEVAN output classified cells based on inferred CNA profiles into likely malignant or normal categories.

### Generation of single-cell reference matrix

A single-cell RNA-seq reference was constructed using our integrated dataset encompassing all cellular populations within the B-ALL niche. A reference expression matrix was generated by extracting raw count data from the RNA assay using GetAssayData() with the “counts” layer. The resulting matrix was converted to a data.frame and matched with corresponding cell-type annotations. The column names of the expression matrix were relabeled with their respective “Annotation_L2” identities to ensure compatibility with CIBERSORT, which requires labeled expression profiles as input. The final reference matrix was saved as a tab-delimited text file (.txt) for use in downstream deconvolution of bulk RNA-seq data.

### Cell type deconvolution analysis in bulk RNA-seq data

Bulk RNA-seq data generated for routine diagnostics were obtained from the Princess Máxima Center Biobank (application number PMCLAB2021.258), containing a cohort of 185 B-ALL bone marrow patient samples. For RNA-seq deconvolution, we employed CIBERSORTx with the Impute Cell Fractions job type, with batch correction disabled, quantile normalization disabled, run mode = absolute, and permutations = 100. Absolute mode scores from CIBERSORTx were used as estimates of relative cell abundance across samples (not direct cell percentages). Resulting Treg estimates were exported to R along with clinical metadata for downstream statistical analysis.

### ERK pathway activity scoring

DESeq2-normalized expression values were obtained from the DESeq2 object (dds), described in ‘Differential expression analysis on bulk RNA seq samples’, using variance-stabilizing transformation. An ERK/MAPK gene set, REACTOME_ERK_MAPK_TARGETS, was retrieved from the MSigDB collection via msigdbr (human species). The gene set was intersected with the rownames of the vst matrix to ensure common identifiers and sets with ≤ 5 matching genes were discarded. Per-sample pathway activity scores were computed by single-sample GSEA (ssGSEA) using the GSVA package. A ssgseaParam object was constructed from the expression matrix and the curated gene set to produce ssGSEA scores (gsva(param)). The resulting matrix (rows = gene set, columns = samples) was transposed to produce one score per sample per gene set and merged with sample metadata and CIBERSORTx Treg estimates. Associations between pathway scores and CIBERSORTx-inferred Treg abundance were assessed using Spearman rank correlation (cor.test, method = “spearman”). All code was run in R (packages: DESeq2, msigdbr, GSVA, dplyr, broom).

### Flow cytometry analysis of T cell phenotypes

Cryopreserved diagnostic B-progenitor ALL bone marrow mononuclear cells were thawed, washed and transferred to a 50 mL tube containing 15 mL complete medium (RPMI-1640 + 10% FBS + 1% penicillin/streptomycin). Cell suspensions were filtered through a 70 µm mesh, counted, centrifuged (350 × g, 5 min) and resuspended at 1 × 10^6 cells/mL. For spectral flow staining, 10 × 10^6 cells were allocated for the full stain and 300,000 cells for an unstained control. Cells were pelleted (400 × g, 5 min) and blocked for Fc receptors with Human TruStain FcX in 1% FBS/PBS for 5 minutes at room temperature. Surface antibody cocktails were added (2.5 µL antibody per 1 × 10^6 cells), incubated 20 minutes protected from light, then washed with 1% FBS/PBS (150 µL) and pelleted (400 × g, 5 min). For intranuclear FOXP3 staining, cells were fixed and permeabilized using the True-Nuclear kit. Cells were stained with the FOXP3-containing antibody cocktail in 50 µL Perm buffer for 20 minutes protected from light, washed three times with Perm buffer, resuspended in 200 µL 1% FBS/PBS, transferred through a filter cap into FACS tubes, brought to volume with an additional 800 µL 1% FBS/PBS, vortexed and acquired on the Cytek Aurora spectral flow cytometer. Detailed antibody clones are provided in Supplementary Table 8.

### T cell proliferation assay

Thawed diagnostic B-ALL BMMCs were washed and resuspended in complete medium (RPMI-1640 + 10% FBS + 1% pen/strep) and counted. Cells were labeled with CFSE (Cell Division Tracker kit, Biolegend, Catalogue 423801) prior to plating. Briefly, CFSE stock was prepared by reconstituting one vial of lyophilized dye in 36 µL DMSO (5 mM). A 5 µM working solution was made by diluting 1 µL of 5 mM stock into 1 mL PBS (1:1,000); working concentration was chosen based on titration. Cells were pelleted and resuspended at 10–100 × 10^6 cells/mL in the CFSE working solution, incubated 20 minutes at room temperature (protected from light), then quenched by adding 5 volumes of complete medium (10% FBS). Cells were pelleted, resuspended in pre-warmed medium and incubated a further 10 minutes at 37°C before final washing. An unstained aliquot was retained as control. Labeled cells were adjusted to 2.5 × 10^6 cells/mL to yield 5 × 10^5 cells per well (200 µL) in sterile U-bottom 96-well plates. Each sample was plated in technical triplicate under two conditions: (1) control medium and (2) stimulation with CD3/CD28 activation beads (Biolegend, Catalogue 422603, 1:1 beads: bone marrow mononuclear cells ratio). Plates were incubated at 37°C/5% CO2. After 5 days, plates were centrifuged (500 × g, 5 min), supernatant removed, and cells stained for surface markers (CD3, isotype control) together with Zombie NIR viability dye for 20 min at room temperature in the dark. Cells were washed, resuspended in 1% FBS/PBS, and acquired by flow cytometry. Data were gated on lymphocytes → singlets → viable CD3+ cells and CFSE dilution was used to assess proliferation. Detailed antibody clones are provided in Supplementary Table 8.

### *Ex vivo* leukemic cell killing assay

Cryopreserved diagnostic B-progenitor ALL bone marrow mononuclear cells (BMMNCs) were thawed, washed and resuspended in complete medium (RPMI 1640 + 10% FBS + 1% pen/strep) at 1 × 10^6 cells/mL. Cells were plated in sterile V-bottom 96-well plates in triplicate (200 µL/well) for four conditions: human IgG control, blinatumomab (1 nM), ipilimumab (100 µg/mL), and blinatumomab (1 nM) + ipilimumab (100 µg/mL). Plates were incubated at 37°C/5% CO2 for 48 hours. At day 2, one well per sample was heat-killed (75°C, 15 min) for live/dead control; remaining wells were centrifuged at 500g, 5 min at 4°C, stained with viability dyes (Annexin V, Viakrome 808) and antibody mix, washed, resuspended in 1% FBS PBS and acquired by flow cytometry (acquisition rate 60 µL/min, recording 40 µL). Data were gated on lymphocytes/ singlets/ CD19+ viable cells (Annexin V–, Viakrome 808–). For each patient, the mean viable CD19+ cell count in IgG control wells was used to normalize treatment wells (mean control set to 1); percent specific leukemic-cell killing was calculated from normalized values.

### Statistical analysis

Unless otherwise stated, statistical significance was assessed by a two-tailed Student’s t-test for comparison between two groups or two-way ANOVA with Bonferroni’s multiple comparison for comparisons among multiple groups. A p value < 0.05 was considered statistically significant. Statistical analyses were performed using R. GSEA is performed with the hypergeometric test for overrepresentation (one-sided), followed by multiple comparison correction using the Benjamini-Hochberg method. The individual method of correction for specific R packages is described in the corresponding publications.

## Data availability

The single-cell RNA sequencing data generated in this study have been deposited in Zenodo at (To be added after acceptance). All other relevant data supporting the key findings of this study are available within the article and its Supplementary Information files or from the corresponding author upon request.

## Code availability

Custom scripts are available at: https://github.com/ (To be added after acceptance) Any additional information required to reanalyze the data reported in this paper is available from the corresponding author upon request.

## Author contributions

M.N.F.B. conceptualized the study, conducted bioinformatic analyses, performed ex vivo experiments, interpreted the data, and drafted and prepared the manuscript. B.K. conducted the ex vivo assays presented in Figure 5 and contributed to manuscript writing and preparation. A.P. conducted the ex vivo assays presented in Figure 1. L.K. performed mutation calling of the bulk-RNA-seq data on our patient cohort. O.H. contributed to supervision and manuscript preparation. H.J.V. oversaw the design, interpretation, and analysis of all experiments and contributed to manuscript writing and preparation.

## Acknowledgements

We thank the single cell sequencing facility and the flow cytometry facility at the Princess Máxima Center for their technical support. We also acknowledge Farnaz Barneh, Babette Hoen, Cesca van de Ven and Monique den Boer, for their advice and contributions.

## Funding

This work has been supported by Horizon Europe’s Marie Sklodowska – Curie Actions co-fund project number 101081481 (Butterfly).

## Declaration of interests

The authors declare no competing interests.

## Notes

### Competing Interest Statement

The authors have declared no competing interest.

## Bibliography

1. Inaba, H. and C.G. Mullighan, Pediatric acute lymphoblastic leukemia. Haematologica, 2020. 105(11): p. 2524–2539.

2. Case, M., E. Matheson, L. Minto, R. Hassan, C.J. Harrison, N. Bown, et al., Mutation of genes affecting the RAS pathway is common in childhood acute lymphoblastic leukemia. Cancer Res, 2008. 68(16): p. 6803–9.

3. Waanders, E., Z. Gu, S.M. Dobson, Z. Antic, J.C. Crawford, X. Ma, et al., Mutational landscape and patterns of clonal evolution in relapsed pediatric acute lymphoblastic leukemia. Blood Cancer Discov, 2020. 1(1): p. 96–111.

4. Brady, S.W., K.G. Roberts, Z. Gu, L. Shi, S. Pounds, D. Pei, et al., The genomic landscape of pediatric acute lymphoblastic leukemia. Nat Genet, 2022. 54(9): p. 1376–1389.

5. Irving, J., E. Matheson, L. Minto, H. Blair, M. Case, C. Halsey, et al., Ras pathway mutations are prevalent in relapsed childhood acute lymphoblastic leukemia and confer sensitivity to MEK inhibition. Blood, 2014. 124(23): p. 3420–30.

6. Jerchel, I.S., A.Q. Hoogkamer, I.M. Aries, E.M.P. Steeghs, J.M. Boer, N.J.M. Besselink, et al., RAS pathway mutations as a predictive biomarker for treatment adaptation in pediatric B-cell precursor acute lymphoblastic leukemia. Leukemia, 2018. 32(4): p. 931–940.

7. Matheson, E.C., H. Thomas, M. Case, H. Blair, R.K. Jackson, D. Masic, et al., Glucocorticoids and selumetinib are highly synergistic in RAS pathway-mutated childhood acute lymphoblastic leukemia through upregulation of BIM. Haematologica, 2019. 104(9): p. 1804–1811.

8. Huang, L., Z. Guo, F. Wang, and L. Fu, KRAS mutation: from undruggable to druggable in cancer. Signal Transduct Target Ther, 2021. 6(1): p. 386.

9. Molina-Arcas, M. and J. Downward, Exploiting the therapeutic implications of KRAS inhibition on tumor immunity. Cancer Cell, 2024. 42(3): p. 338–357.

10. Tian, J., J.H. Chen, S.X. Chao, K. Pelka, M. Giannakis, J. Hess, et al., Combined PD-1, BRAF and MEK inhibition in BRAF(V600E) colorectal cancer: a phase 2 trial. Nat Med, 2023. 29(2): p. 458–466.

11. Canon, J., K. Rex, A.Y. Saiki, C. Mohr, K. Cooke, D. Bagal, et al., The clinical KRAS(G12C) inhibitor AMG 510 drives anti-tumour immunity. Nature, 2019. 575(7781): p. 217–223.

12. von Stackelberg, A., F. Locatelli, G. Zugmaier, R. Handgretinger, T.M. Trippett, C. Rizzari, et al., Phase I/Phase II Study of Blinatumomab in Pediatric Patients With Relapsed/Refractory Acute Lymphoblastic Leukemia. J Clin Oncol, 2016. 34(36): p. 4381–4389.

13. Bassan, R., S. Chiaretti, I. Della Starza, A. Santoro, O. Spinelli, M. Tosi, et al., Up-front blinatumomab improves MRD clearance and outcome in adult Ph-B-lineage ALL: the GIMEMA LAL2317 phase 2 study. Blood, 2025. 145(21): p. 2447–2459.

14. van der Sluis, I.M., P. de Lorenzo, R.S. Kotecha, A. Attarbaschi, G. Escherich, K. Nysom, et al., Blinatumomab Added to Chemotherapy in Infant Lymphoblastic Leukemia. N Engl J Med, 2023. 388(17): p. 1572–1581.

15. Locatelli, F., G. Zugmaier, C. Rizzari, J.D. Morris, B. Gruhn, T. Klingebiel, et al., Effect of Blinatumomab vs Chemotherapy on Event-Free Survival Among Children With High-risk First-Relapse B-Cell Acute Lymphoblastic Leukemia: A Randomized Clinical Trial. JAMA, 2021. 325(9): p. 843–854.

16. Gupta, S., R.E. Rau, J.A. Kairalla, K.R. Rabin, C. Wang, A.L. Angiolillo, et al., Blinatumomab in Standard-Risk B-Cell Acute Lymphoblastic Leukemia in Children. N Engl J Med, 2025. 392(9): p. 875–891.

17. Kantarjian, H., A. Stein, N. Gokbuget, A.K. Fielding, A.C. Schuh, J.M. Ribera, et al., Blinatumomab versus Chemotherapy for Advanced Acute Lymphoblastic Leukemia. N Engl J Med, 2017. 376(9): p. 836–847.

18. Litzow, M.R., Z. Sun, R.J. Mattison, E.M. Paietta, K.G. Roberts, Y. Zhang, et al., Blinatumomab for MRD-Negative Acute Lymphoblastic Leukemia in Adults. N Engl J Med, 2024. 391(4): p. 320–333.

19. Bendig, S., M. Kelm, A. Laqua, M. Szczepanowski, T. Beder, N. Wolgast, et al., Baseline T-cell fitness and subtype-specific CD19 loss as drivers of blinatumomab resistance in B-ALL. Blood, 2025. 146(Supplement 1): p. 32–32.

20. Ferrao Blanco, M.N., B. Kazybay, M. Belderbos, O. Heidenreich, and H.J. Vormoor, Distinct stromal cell populations define the B-cell acute lymphoblastic leukemia microenvironment. Leukemia, 2025. 39(11): p. 2622–2639.

21. Khan, O., J.R. Giles, S. McDonald, S. Manne, S.F. Ngiow, K.P. Patel, et al., TOX transcriptionally and epigenetically programs CD8(+) T cell exhaustion. Nature, 2019. 571(7764): p. 211–218.

22. Zheng, L., S. Qin, W. Si, A. Wang, B. Xing, R. Gao, et al., Pan-cancer single-cell landscape of tumor-infiltrating T cells. Science, 2021. 374(6574): p. abe6474.

23. Good, C.R., M.A. Aznar, S. Kuramitsu, P. Samareh, S. Agarwal, G. Donahue, et al., An NK-like CAR T cell transition in CAR T cell dysfunction. Cell, 2021. 184(25): p. 6081–6100 e26.

24. Anderson, Ana C., N. Joller, and Vijay K. Kuchroo, Lag-3, Tim-3, and TIGIT: Co-inhibitory Receptors with Specialized Functions in Immune Regulation. Immunity, 2016. 44(5): p. 989–1004.

25. Marelli-Berg, F.M., M. Clement, C. Mauro, and G. Caligiuri, An immunologist’s guide to CD31 function in T-cells. J Cell Sci, 2013. 126(Pt 11): p. 2343–52.

26. Duell, J., M. Dittrich, T. Bedke, T. Mueller, F. Eisele, A. Rosenwald, et al., Frequency of regulatory T cells determines the outcome of the T-cell-engaging antibody blinatumomab in patients with B-precursor ALL. Leukemia, 2017. 31(10): p. 2181–2190.

27. Hoen, B.D.J., T.J.J. Manuputty, J.M. Boer, C. Van De Ven, and M.L. den Boer, The Leukemic Microenvironment Represses Blinatumomab-Dependent Killing in Pediatric B-Cell Precursor Acute Lymphoblastic Leukemia. Blood, 2024. 144(Supplement 1): p. 4228–4228.

28. Wing, K., Y. Onishi, P. Prieto-Martin, T. Yamaguchi, M. Miyara, Z. Fehervari, et al., CTLA-4 control over Foxp3+ regulatory T cell function. Science, 2008. 322(5899): p. 271–5.

29. van Maldegem, F. and J. Downward, Mutant KRAS at the Heart of Tumor Immune Evasion. Immunity, 2020. 52(1): p. 14–16.

30. Yuan, S., K.S. Stewart, Y. Yang, M.D. Abdusselamoglu, S.M. Parigi, T.Y. Feinberg, et al., Ras drives malignancy through stem cell crosstalk with the microenvironment. Nature, 2022. 612(7940): p. 555–563.

31. Wing, K. and S. Sakaguchi, Regulatory T cells exert checks and balances on self tolerance and autoimmunity. Nat Immunol, 2010. 11(1): p. 7–13.

32. Hall, A.G. and R.E. Rau, Blinatumomab use in pediatric B-ALL: where are we now? Blood Adv, 2025. 9(15): p. 3946–3954.

33. Kantarjian, H., I. Aldoss, and E. Jabbour, Management of Adult Acute Lymphoblastic Leukemia: A Review. JAMA Oncol, 2025. 11(7): p. 771–778.

34. Patricelli, M.P., M.R. Janes, L.S. Li, R. Hansen, U. Peters, L.V. Kessler, et al., Selective Inhibition of Oncogenic KRAS Output with Small Molecules Targeting the Inactive State. Cancer Discov, 2016. 6(3): p. 316–29.

35. Hong, D.S., M.G. Fakih, J.H. Strickler, J. Desai, G.A. Durm, G.I. Shapiro, et al., KRAS(G12C) Inhibition with Sotorasib in Advanced Solid Tumors. N Engl J Med, 2020. 383(13): p. 1207–1217.

36. Menne, T., D. Slade, J. Savage, S. Johnson, J. Irving, P. Kearns, et al., Selumetinib in combination with dexamethasone for the treatment of relapsed/refractory RAS-pathway mutated paediatric and adult acute lymphoblastic leukaemia (SeluDex): study protocol for an international, parallel-group, dose-finding with expansion phase I/II trial. BMJ Open, 2022. 12(3): p. e059872.

37. Vormoor, B., J. Savage, C. Kristunas, S. Johnson, M.N.F. Blanco, G. Shenton, et al., Responses can be achieved with a combination of the MEK inhibitor selumetinib and dexamethasone in patients with relapsed/refractory RAS-pathway mutated acute lymphoblastic leukemia: Results of a parallel cohort, dose-finding and expansion phase I/II trial. EJC Paediatric Oncology, 2026. 7.

38. Love, M.I., W. Huber, and S. Anders, Moderated estimation of fold change and dispersion for RNA-seq data with DESeq2. Genome Biol, 2014. 15(12): p. 550.

39. Germain, P.L., A. Lun, C. Garcia Meixide, W. Macnair, and M.D. Robinson, Doublet identification in single-cell sequencing data using scDblFinder. F1000Res, 2021. 10: p. 979.

40. Bandyopadhyay, S., M.P. Duffy, K.J. Ahn, J.H. Sussman, M. Pang, D. Smith, et al., Mapping the cellular biogeography of human bone marrow niches using single-cell transcriptomics and proteomic imaging. Cell, 2024. 187(12): p. 3120–3140 e29.

41. Stuart, T., A. Butler, P. Hoffman, C. Hafemeister, E. Papalexi, W.M. Mauck, 3rd, et al., Comprehensive Integration of Single-Cell Data. Cell, 2019. 177(7): p. 1888–1902 e21.

42. Aran, D., A.P. Looney, L. Liu, E. Wu, V. Fong, A. Hsu, et al., Reference-based analysis of lung single-cell sequencing reveals a transitional profibrotic macrophage. Nat Immunol, 2019. 20(2): p. 163–172.

43. Jin, S., C.F. Guerrero-Juarez, L. Zhang, I. Chang, R. Ramos, C.H. Kuan, et al., Inference and analysis of cell-cell communication using CellChat. Nat Commun, 2021. 12(1): p. 1088.

44. De Falco, A., F. Caruso, X.D. Su, A. Iavarone, and M. Ceccarelli, A variational algorithm to detect the clonal copy number substructure of tumors from scRNA-seq data. Nat Commun, 2023. 14(1): p. 1074.

