## Supplementary_Figures for "The bone marrow microenvironment of RAS pathway mutant B-ALL is enriched for immunosuppressive regulatory T cells"

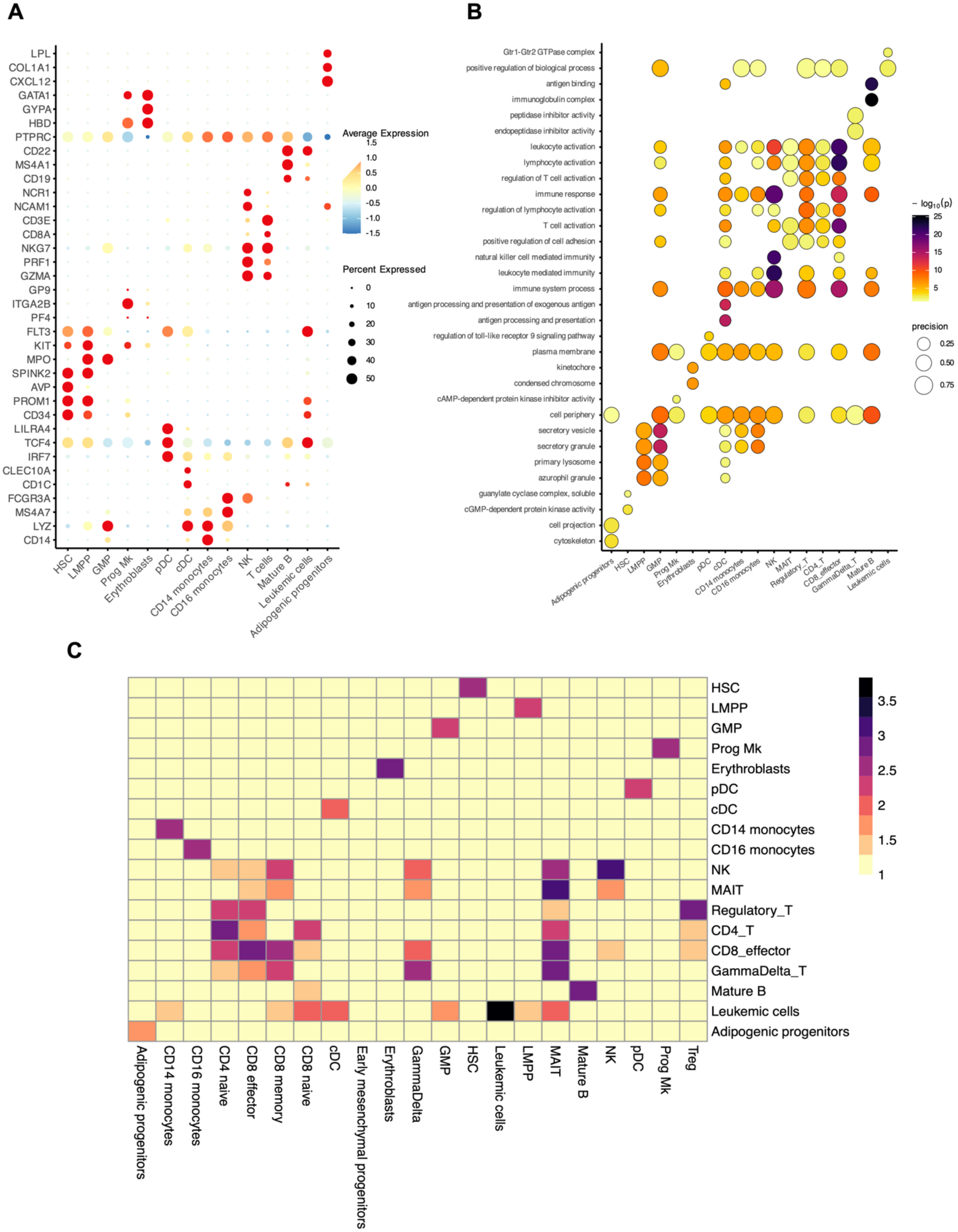

Supplementary Figure 1. A) Dot plot of canonical marker genes across annotated cell populations. Dot size indicates percent of cells expressing the gene per cluster; color indicates scaled average expression. Markers include lineage and functional genes used to annotate major cell types. B) Cluster-resolved gene-ontology enrichment summary. Top enriched GO/REAC/KEGG terms for upregulated marker genes per cluster (top 2 shown). Plots display GO terms most strongly associated with each cluster. C) Heatmap comparing the cell type annotation of the single cell dataset in this study (y-axis) to the reference labels (x-axis, Ferrao Blanco et al., Leukemia 2025). Values are log10(count+10) of cell counts per annotation pair. Heatmap shows correspondence between annotated clusters and reference classes.

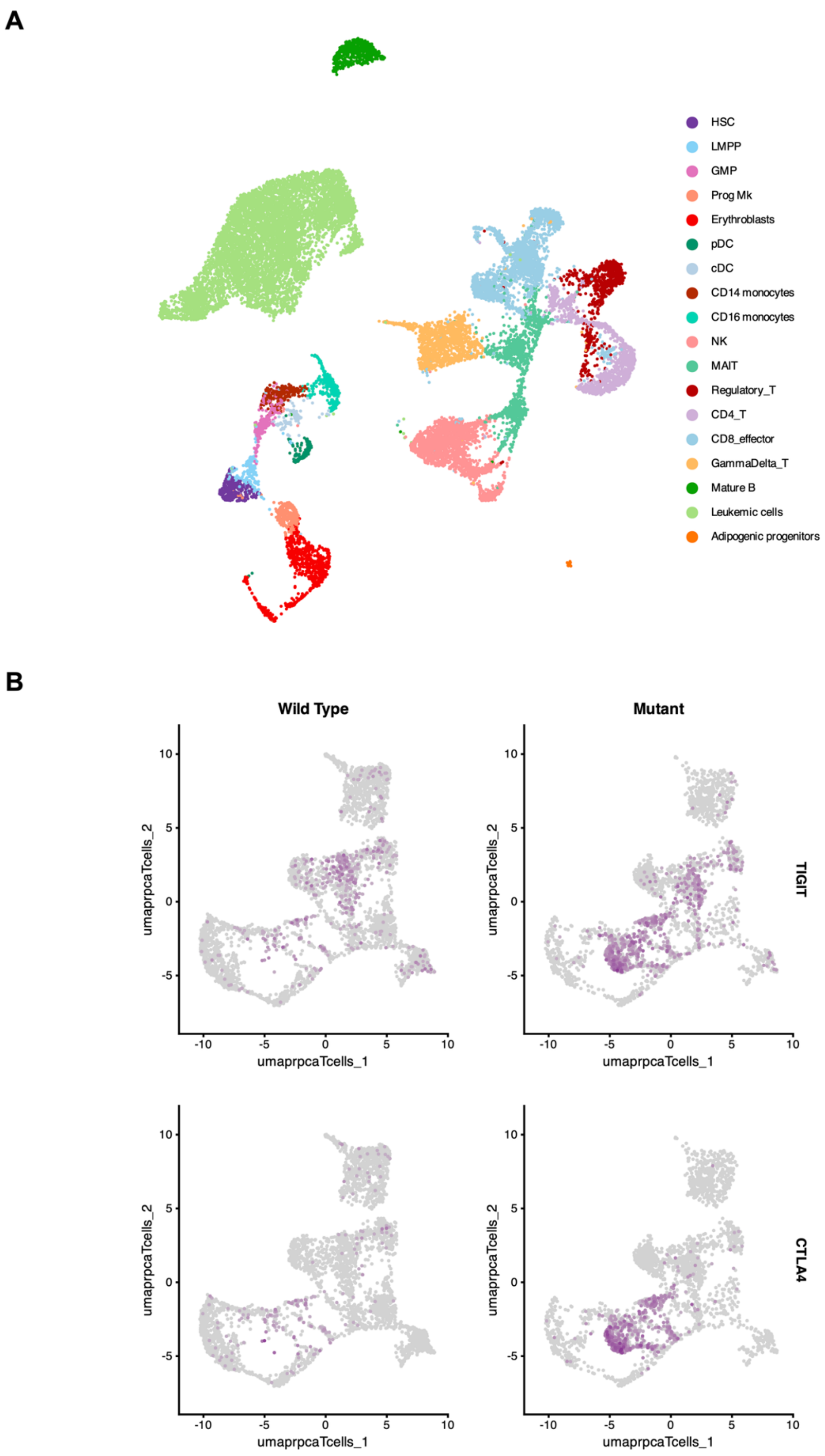
A
HSC
LMPP
GMP
Prog Mk
Erythroblasts
pDC
cDC
CD14 monocytes
CD16 monocytes
NK
MAIT
Regulatory_T
CD4_T
CD8_effector
GammaDelta_T
Mature B
Leukemic cells
Adipogenic progenitors
B
Wild Type
Mutant
TIGIT
CTLA4
umaprpcaTcells_1
umaprpcaTcells_2
10
5
0
-5
-10

Supplementary Figure 2. A) UMAP of bone marrow mononuclear cells showing refined annotation of T-cell populations (labels indicated). B) Feature plots of TIGIT and CTLA4 expression across the T-cell compartment (UMAP projection as in Figure 2D).

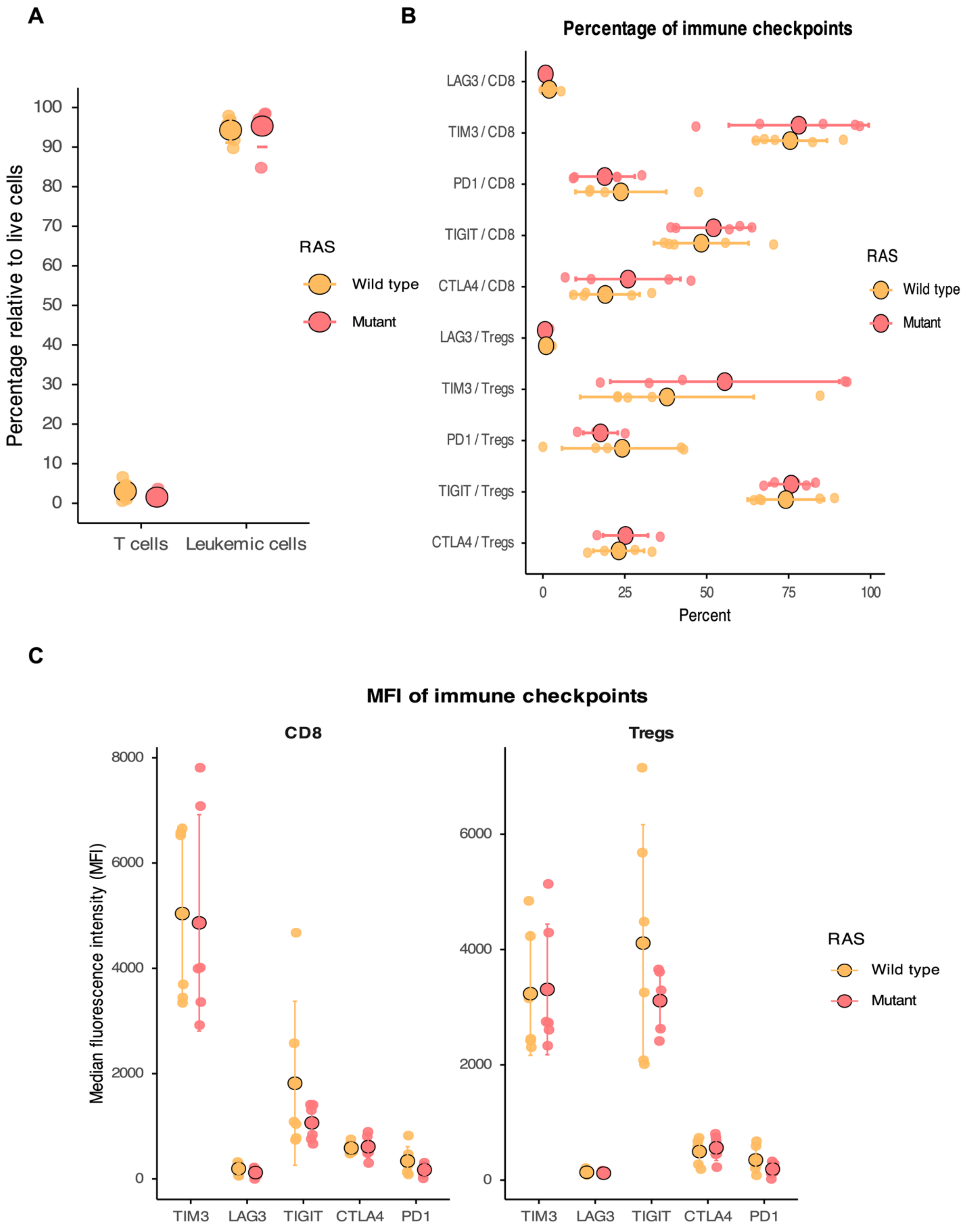
A
Percentage relative to live cells
100
90
80
70
60
50
40
30
20
10
0
T cells
Leukemic cells
RAS
Wild type
Mutant
B
Percentage of immune checkpoints
LAG3 / CD8
TIM3 / CD8
PD1 / CD8
TIGIT / CD8
CTLA4 / CD8
LAG3 / Tregs
TIM3 / Tregs
PD1 / Tregs
TIGIT / Tregs
CTLA4 / Tregs
0
25
50
75
100
Percent
RAS
Wild type
Mutant
C
MFI of immune checkpoints
CD8
Tregs
Median fluorescence intensity (MFI)
8000
6000
4000
2000
0
TIM3
LAG3
TIGIT
CTLA4
PD1
RAS
Wild type
Mutant

Supplementary Figure 3. A) Spectral flow cytometry analysis of leukemic cells and total CD3+ T cells in live bone marrow cells (n = 12; 6 RAS-mutant, 6 wild-type samples). B) Frequency (%) of checkpoint-positive cells in regulatory T cells (Tregs) and CD8+ T cells measured by spectral flow cytometry (same cohort). C) Median fluorescence intensity (MFI) of checkpoint/exhaustion markers in T cell subsets from bone marrow (same cohort). Statistical comparisons were performed with Welch's t-test and Bonferroni correction for multiple comparisons. No significant differences were observed between groups for all comparisons shown in panels A, B and C.

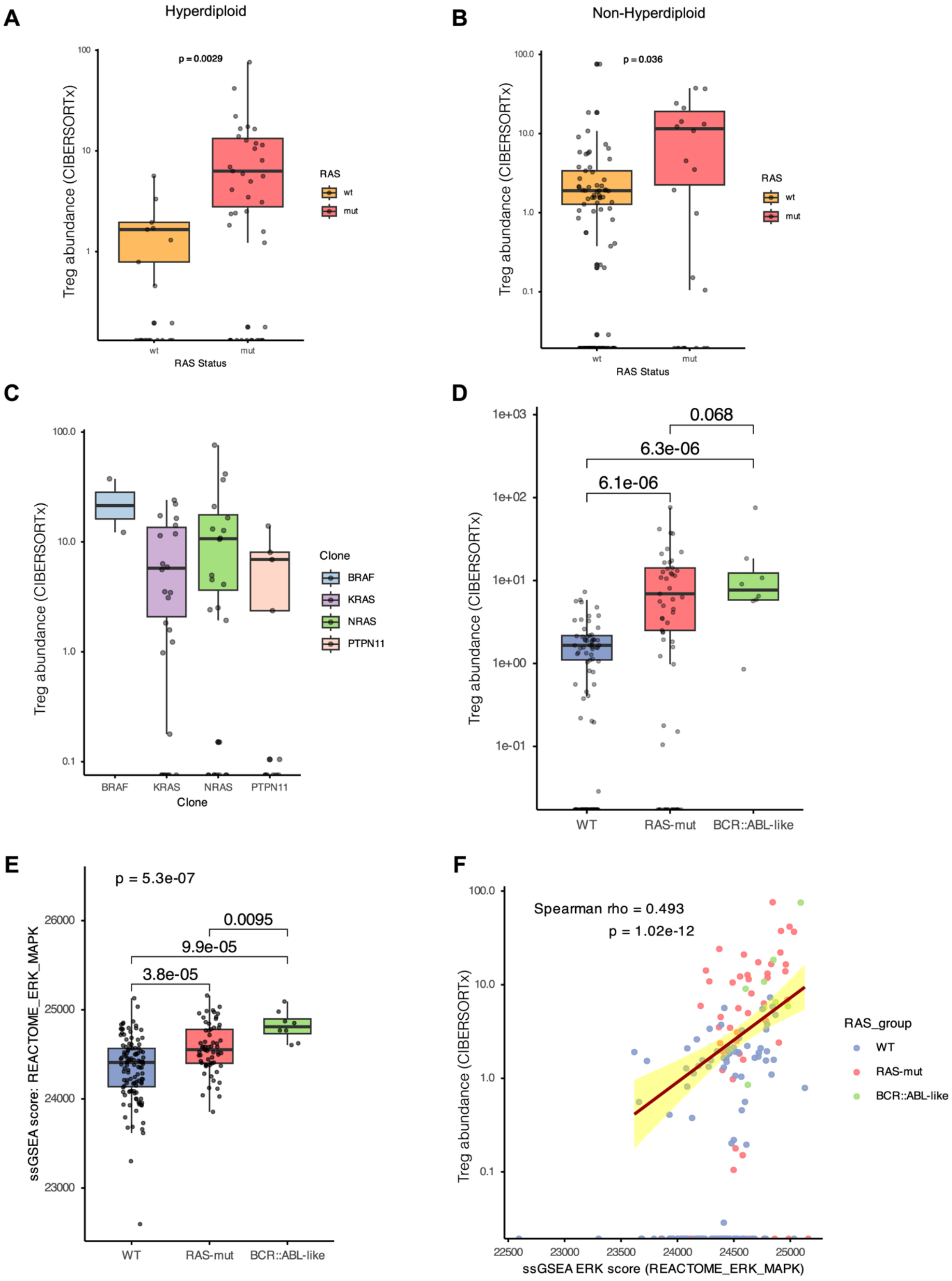
A
Hyperdiploid
p = 0.0029
Treg abundance (CIBERSORTx)
RAS Status
wt
mut
RAS
B
Non-Hyperdiploid
p = 0.036
C
Clone
BRAF
KRAS
NRAS
PTPN11
D
0.068
6.3e-06
6.1e-06
WT
RAS-mut
BCR::ABL-like
E
p = 5.3e-07
0.0095
9.9e-05
3.8e-05
ssGSEA score: REACTOME_ERK_MAPK
F
Spearman rho = 0.493
p = 1.02e-12
RAS_group
ssGSEA ERK score (REACTOME_ERK_MAPK)

Supplementary Figure 4. CIBERSORTx deconvolution of bulk RNA-seq (n = 185) and ERK pathway analysis. A) CIBERSORTx estimates of regulatory T cell abundance (absolute scores) restricted to hyperdiploid cases. Group comparison (RAS-mutant vs wild-type) was performed by two-sided Student's t-test. B) CIBERSORTx estimates of regulatory T cells in combined non-hyperdiploid subtypes; two-sided Student's t-test was used for group comparison. C) CIBERSORTx Treg estimates stratified by specific RAS-pathway mutated gene (KRAS, NRAS, PTPN11, BRAF). Differences across mutation groups were tested by one-way ANOVA with Tukey's HSD post-hoc pairwise comparisons; no significant differences were observed. D) CIBERSORTx Treg abundance comparing wild-type, RAS-mutant and BCR::ABL (-like) samples (combined). Differences were tested by Kruskal–Wallis; pairwise comparisons were performed with Wilcoxon rank-sum tests with Bonferroni correction. Exact p-values are reported on the panels. E) Boxplot of per-sample ssGSEA scores for the REACTOME_ERK_MAPK gene set by RAS_group (WT, RAS-mut, BCR::ABL-like). Group differences were assessed by Kruskal–Wallis test with Bonferroni-corrected pairwise Wilcoxon tests; pairwise comparisons are annotated on the plot. F) Scatterplot of REACTOME_ERK_MAPK ssGSEA score versus CIBERSORTx Treg absolute score (log10 scaled on the y-axis). Points are colored by RAS_group; a single overall linear regression line (±95% CI) is shown. Spearman correlation (rho and p-value) between REACTOME_ERK_MAPK and log10(1 + Treg score) is reported on the panel.
